# 3D printing and bioprinting for miniaturized and scalable hanging-drop organoids culture

**DOI:** 10.1101/2025.09.25.678315

**Authors:** Elena Bianchi, Oronza A. Botrugno, Paola De Stefano, Giovanni F. M. Gallo, Claudia Felici, Jolanda M. Bruno, Giulio Giovannoni, Francesca Ratti, Luca A. Aldrighetti, Roger D. Kamm, Giovanni Tonon, Gabriele Dubini

## Abstract

Three-dimensional (3D) cell culture systems rely on the manipulation of a biologically derived matrix, typically soluble Basement Membrane Extract (sBME), in which cells or cellular aggregates, such as organoids, are suspended. This matrix provides mechanobiological support, promoting cellular processes. However, the handling of sBME-based matrices containing cellular constructs poses significant challenges due to their rheological properties. We developed an integrated bioprinting system to surpass the conventional pipetting, seeding and culture in multiwell plates. The system combines a fluidic cartridge with innovative 3D-printed biocompatible culture tools designed to host and preserve high-throughput microcultures of Patient-Derived Organoids (PDOs) in sBME. The miniaturized hanging-drop configuration enables extended culture periods and high-throughput imaging screenings. This comprehensive approach overcomes common issues associated with sBME, including sedimentation of cellular aggregates, premature gelation, and structural collapse, which negatively impact culture quality and reproducibility throughout the entire 3D culture workflow, from seeding to culture maintenance, and post-culture analyses.

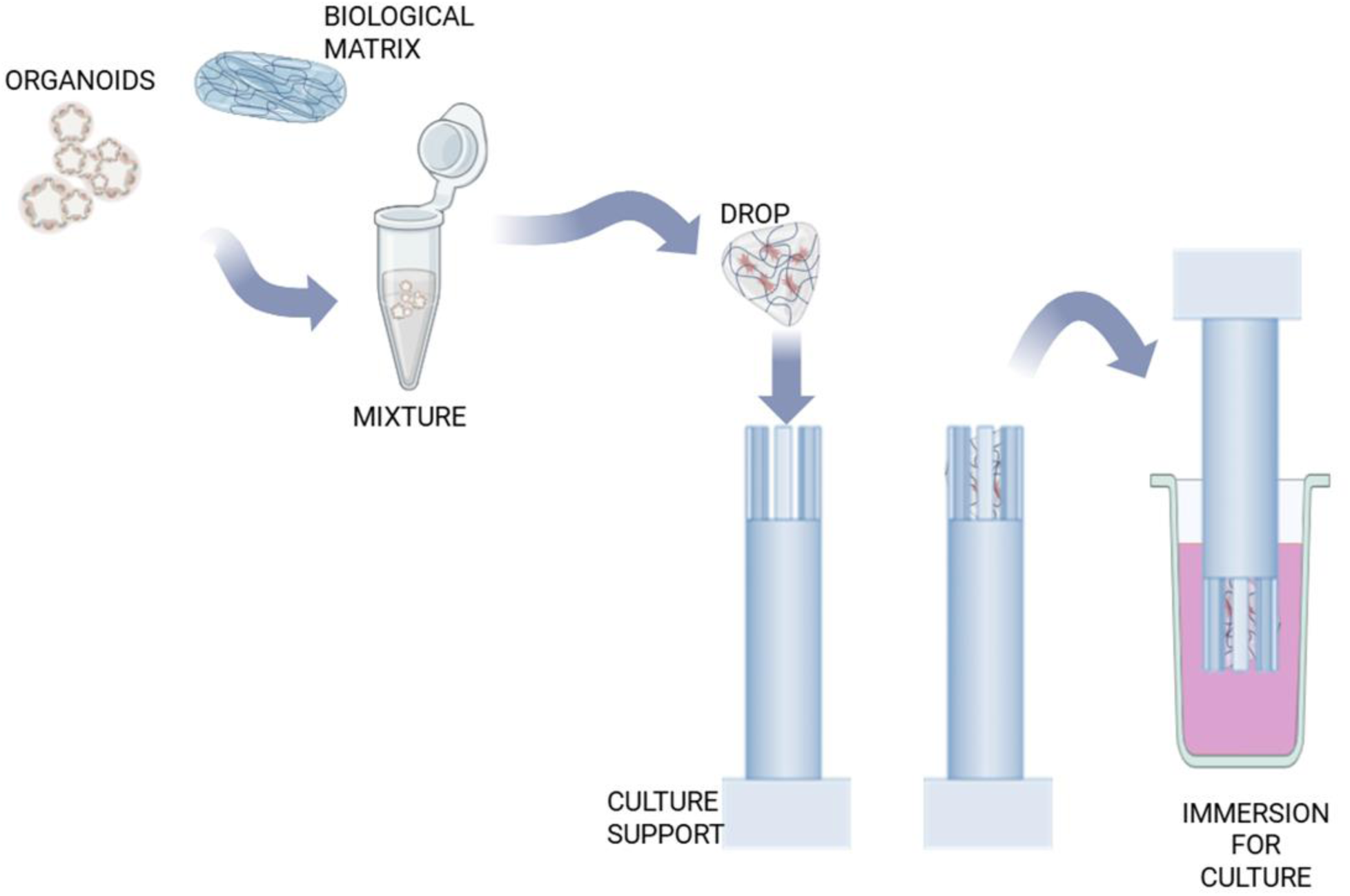

**Highlights:**

- Miniaturized 3D hanging-drop matrix-embedded organoid culture in a 384-well plate
- Custom cartridge enables homogeneous bioprinting of organoids in sBME-based matrix
- 3D-printed tools support compact, scalable multiwell culture systems
- System suited for miniaturized culture organoids for high-throughput drug screening
- Scalable miniaturized culture system for extended periods of time

---

Organoids are self-organizing organotypic three-dimensional (3D) structures that can be derived from single stem cells and generate different cell populations through proliferation and differentiation, recapitulating the structural and functional organization found in the tissue of origin ^1^. Unlike two-dimensional (2D) cultures, organoids provide sophisticated physiological and pathophysiological models, offering improved consistency and robustness towards in vitro experimental outcomes ^2–4^. A 3D support structure, usually in the form of a hydrogel scaffold, is an essential component of organoids culture. The topological, mechanical, and biochemical features of this structure are crucial for the development of the culture ^5^. The hydrogel provides biomechanical support and enables cells to self-assemble into complex architectures that mimic the intrinsic organization of their tissue of origin ^6^. It provides nutrients and allows the permeation of both the culture medium and oxygen. Matrigel™ (Corning, Corning, NY, USA) is considered the “gold standard” among the extracellular matrices (ECMs). It is a soluble basement membrane extract (sBME) derived from Engelbreth-Holm-Swarm (EHS) mouse sarcoma and is primarily composed of laminin and collagen type IV ^7^. Alternative products, such as Cultrex™ (R&D Systems, Minneapolis, USA), contain a similar heterogeneous mixture of proteins and growth factors ^8^.

Patient-derived organoids (PDOs) are transforming biomedical research ^5^, as they retain cellular heterogeneity while faithfully mimicking the histopathological and genomic characteristics of their tissue of origin. A strong concordance between drug responses in patients and their corresponding PDOs has been demonstrated across multiple tumor types ^9^, reinforcing their potential as predictive avatars for personalized therapy ^10^, especially in cancer research ^11^. The feasibility of medium-throughput drug screenings on PDOs has been demonstrated by several groups ^12^, but several constraints, including the incorporation of multiple cell types, the heterogeneity in size, volume and density, and the embedding in ECM, have limited their full application in high-throughput drug screenings. Handling sBMEs in their pure form remains very challenging when using pipettes, dispensing robots or pneumatically driven bioprinters ^13,14^ due to their complex non-Newtonian rheology, irreversible thermal gelation, and low mechanical properties. These features have, so far, prevented their widespread use in high-throughput screening (HTS) assays, which rely on liquid-handling robots. When prepared for drug screening, ECM is removed and organoids are suspended in media at a low percentage of sBME ^4,15–17^. In bioprinting, the rheology of ECM limits direct pressure-controlled extrusion, prompting the use of blended bioinks or syringe-based dispensers with volumetric control. Extruding biological matrices with organoid aggregates can result in uneven seeding due to sedimentation ^18^. Although solutions such as sequential bioprinting ^19^ or embedding in pre-polymerized matrices ^13^ address compartmentalization ^20^, they are not compatible with microfluidic samples. In microfluidic systems, micropillars and phase guides can help to localize and retain constructs ^21–25^; however, conventional sealed chips can make seeding and retrieval difficult, which limits laboratory integration ^26,27^.

There is a pressing need to develop strategies that improve the reproducibility, miniaturization, and scalability of organoid seeding and culture in extracellular sBME-based matrix substitutes ^10^, particularly to enable the use of high-concentration sBME matrices in miniaturized formats for high-throughput drug screening by addressing challenges related to rheology of the matrices, constructs sedimentation, and culture compartmentalization.

We developed a miniaturized culture system consisting of 3D-printed structures, fabricated from biocompatible resin, which overcomes these limitations related to rheology, construct sedimentation, and compartmentalization. These structures are designed to host and preserve intact miniaturized organoid cultures over extended periods of time. Additionally, we implemented a dispensing system capable of bioprinting microliter-scale droplets of a homogeneous matrix– organoid suspension into custom 3D-printed structures.

## Results

### Development of a bioprinting system for seeding of organoids embedded in a matrix in a miniaturized culture format

We have developed a seeding system, consisting of a custom-designed cartridge mounted on a volumetric pump (Fig. 1). This system enables the consistent and uniform dispensing of droplets of a suspension of biologically derived matrices based on sBME and organoids. These droplets are successfully bioprinted into the NESTs, microstructures fabricated via 3D printing using a biocompatible resin. The structures are sized to accommodate volumes of 4, 6, and 8 µL, respectively. Since the flow rate for the extrusion of the drops is set at 200 μL/min, the extrusion and settling times are proportional to the extruded volumes: 1,2, 1,8 and 2,4 s, respectively, for 4, 6, and 8 µL. Upon extrusion from the cartridge, each droplet adheres to the target structure and fully transfers into the internal cavity and fills it (Fig. 2a). Throughout both the seeding and subsequent gelation phases, the droplet remains stably anchored to the cavity, even when the structure is inverted and placed in culture. We did not observe any structural collapse of the droplets, including during the gelation step. Since the extrusion is performed by a commercial high-accuracy syringe pump, the extruded volumes are consistent. Indeed, to assess whether the organoids seeding was homogeneous, we measured ATP content in the dispensed droplets immediately after seeding. This value was used as a proxy for the number of seeded organoids. Since the estimated concentration of organoids is the same for all the volumes in this assay, the luminescence values proportionally increased with the volume of the droplets fitting a linear trend (Fig. 2b). Moreover, the homogeneity of seeding, in terms of concentration of organoids suspended in the seeded sample, was not affected by the concentration or the type of matrix used to embed organoids (Figs. 2-4 and Fig. 1S). These data contribute to demonstrating that our system enables homogeneous seeding of organoids in a miniaturized format.

**Figure 1.**
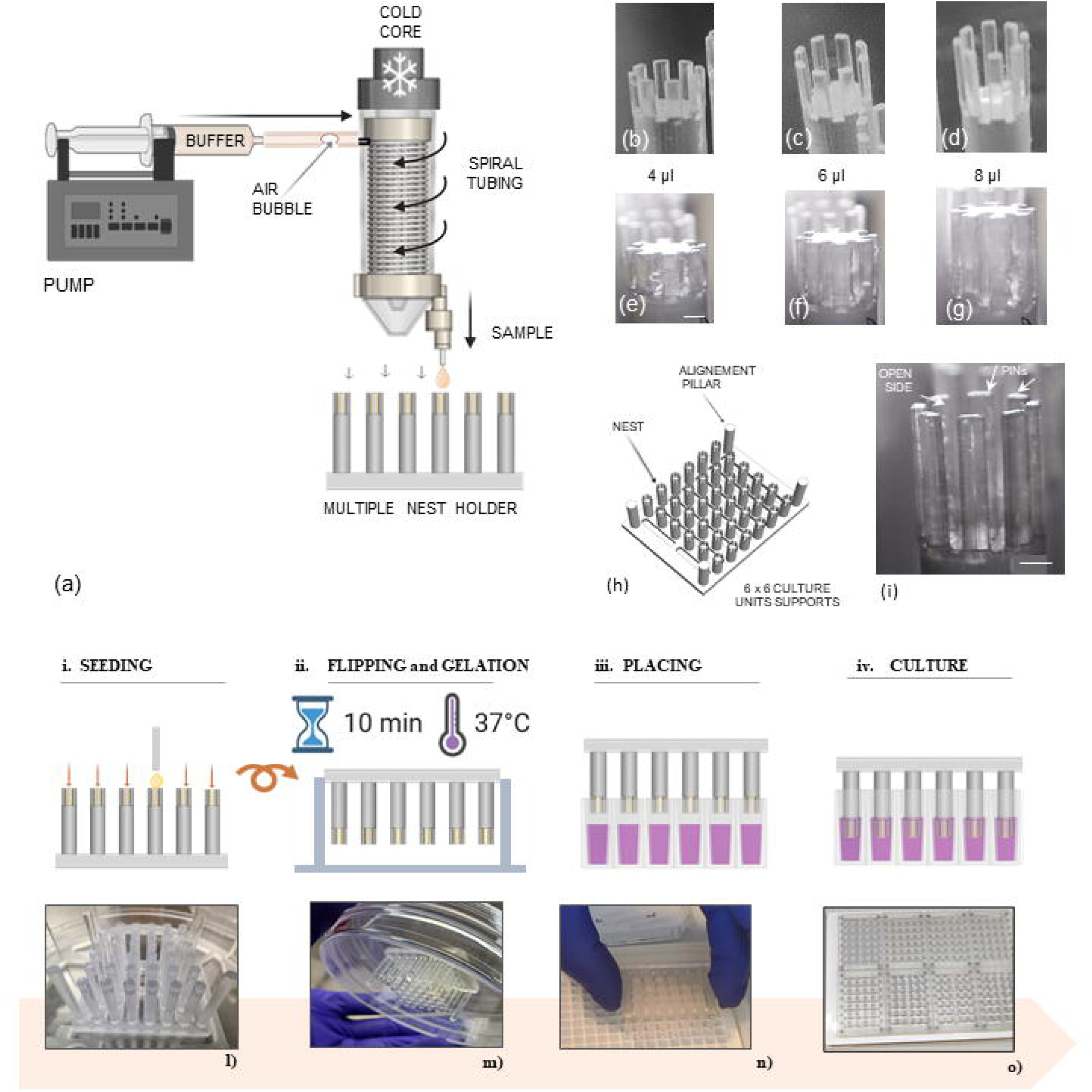
Components of the culture system and stepwise operational procedures of the seeding protocol. a) Schematic representation of the culture system: a pump to activate the extrusion of the buffer (PBS); an air bubble to avoid mixing of the sample and PBS; the spiral tube consisting of a coil containing the suspension of organoids in matrix, wrapped around a cold core; the extrusion of drops of the suspension of organoids into the NESTs. b), c), d) 3D printed empty NESTs of 4, 6, 8 µL capacities, respectively, in the seeding position (with the open side at the top). e) f) g) NESTs filled with 4, 6, 8 µL volumes, respectively, of 5 mg/ml Matrigel^TM^. h) schematic representation of a 36-unit array of NESTs 3D-printed on a unique holder. On the corners, the four alignment pillars allow the NESTs to be centered in each well and at a fixed distance from the bottom. i) A single NEST with the top open side and the pins arranged all around the circumference of the culture unit alternating with open fenestrations. Scale bar: 0,5 mm. The seeding protocol is defined in four steps: i) sequential seeding of the suspensions of organoids in matrix into the NESTs; ii) overturning of the 36-NESTs holder locked inside a closed Petri dish by a customized 3D-printed tool; iii) manual alignment of the supports into a 384-well plate iv) insertion of the NESTs into the wells filled with medium. l)- o) Representative images of the operational procedures described above. l) The 6x6 culture unit support is filled with matrix-embedded organoids. m) The support is turned upside down during the gelation time interval. n) The 36-NESTs holder is placed in immersion in a 384-well plate. o) Configuration of a 384-well plate setup consisting of 8 pieces of 36-NESTs holders organized for organoids culture.

**Figure 2.**
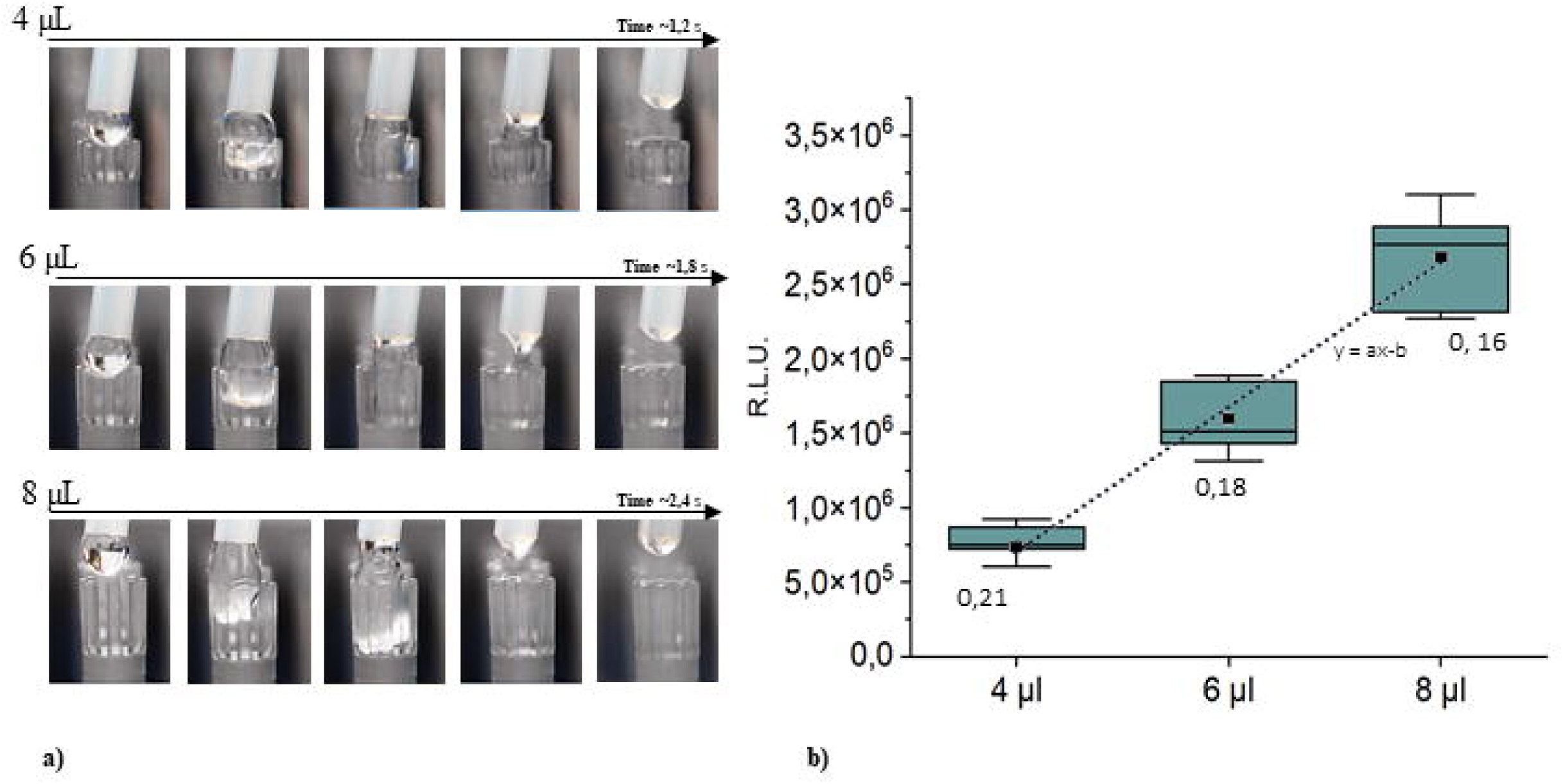
The system allows homogenous seeding of Matrigel^TM^-embedded organoids in drops of different volumes. The homogeneity of seeding of Matrigel-embedded organoids across drops of varying volumes (4, 6, 8 µL) has been tested and validated. a) The time frames report the process of seeding the drops in specific NESTs supports. The flow rate was set to 200 µL/min. The process of seeding involves the initial extrusion of the drop, which comes into contact with the structures, before the drop settles inside the nest and detaches from the tube. Videos used as the source for the screenshots were acquired with the Microqubic imaging system (MRCL700 3D Imager Pro, Mircoqubic AG, Zug, CH). b) PDO#17 suspended in 5mg/ml Matrigel was seeded as reported in a). Organoids were left to recover, and ATP content was measured by CellTiter-Glo assay 1 hour after seeding. Luminescence values are reported as R.L.U. Each condition consists of n=12 replicates. The box chart is defined by 25%-75% of the distribution ± SD (Standard Deviation). The coefficient of variation (CV) is reported for each group. Linear trend of the means a = 4,9·10 ^5^ b =1,2 10 ^6^ R^2^=0,996 .

**Figure 3.**
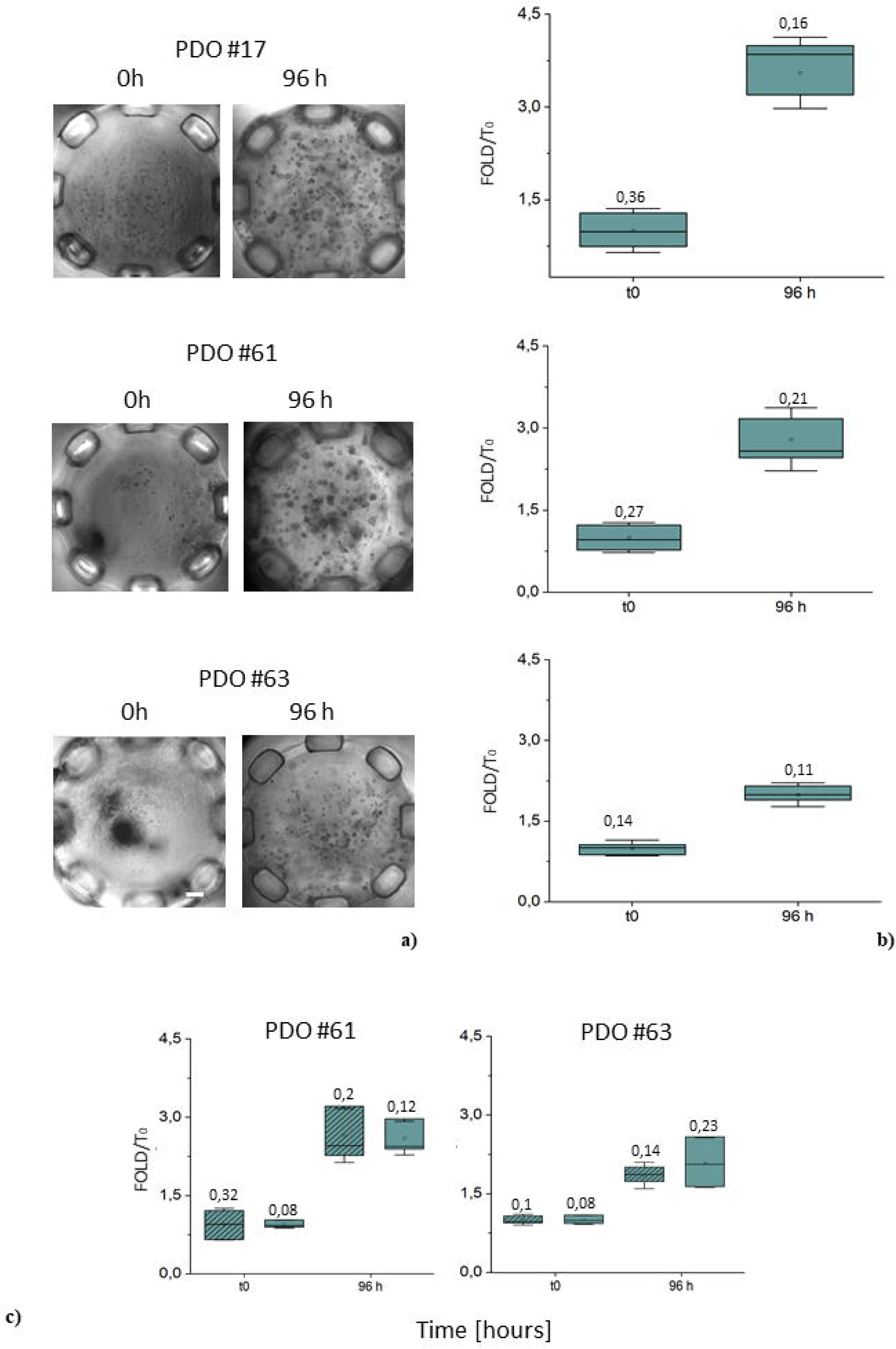
Seeding and culturing of Matrigel^TM^-embedded organoids in a standard 96-well plate and in the NESTs system. a) and b) PDO#17, PDO#61 and PDO#63, suspended in 5mg/ml Matrigel, were seeded (t0) and cultured in the NESTs for 96 hours. a) Representative brightfield images examples, 5X objective lens, of the organoids at each time point. Scale bar: 200 µm. b) The ATP content of organoids was measured by CellTiter-Glo at each time point. n=12 replicates. Growth rate (fold/t0) is reported as the ratio of the luminescence values at each time point relative to that at t0. The box chart is defined by 25%-75% of the distribution ± SD (Standard Deviation). The square symbol represents the average, while the line represents the median value. The coefficient of variation CV is reported for each group. c) PDO#61 and PDO#63, suspended in 5mg/ml Matrigel^TM^ were seeded and cultured in a 96-well plate for 96 hours. In the graph, comparison of the growth rates in the 96-well (solid color) and in the NESTs (stripes pattern) setting. The ATP content of organoids was measured by CellTiter-Glo at the time of seeding (t0) and after 96 hours of culture. For each time point, replicates were n=3 in the 96-well and n=6 in the NESTs system. Growth rate (fold/t0) is reported as the ratio of the luminescence values at each time point relative to that at t0. The box chart is defined by 25%-75% of the distribution ± SD (Standard Deviation). The square symbol represents the average, while the line represents the median value. The coefficient of variation CV is reported for each group.

**Figure 4.**
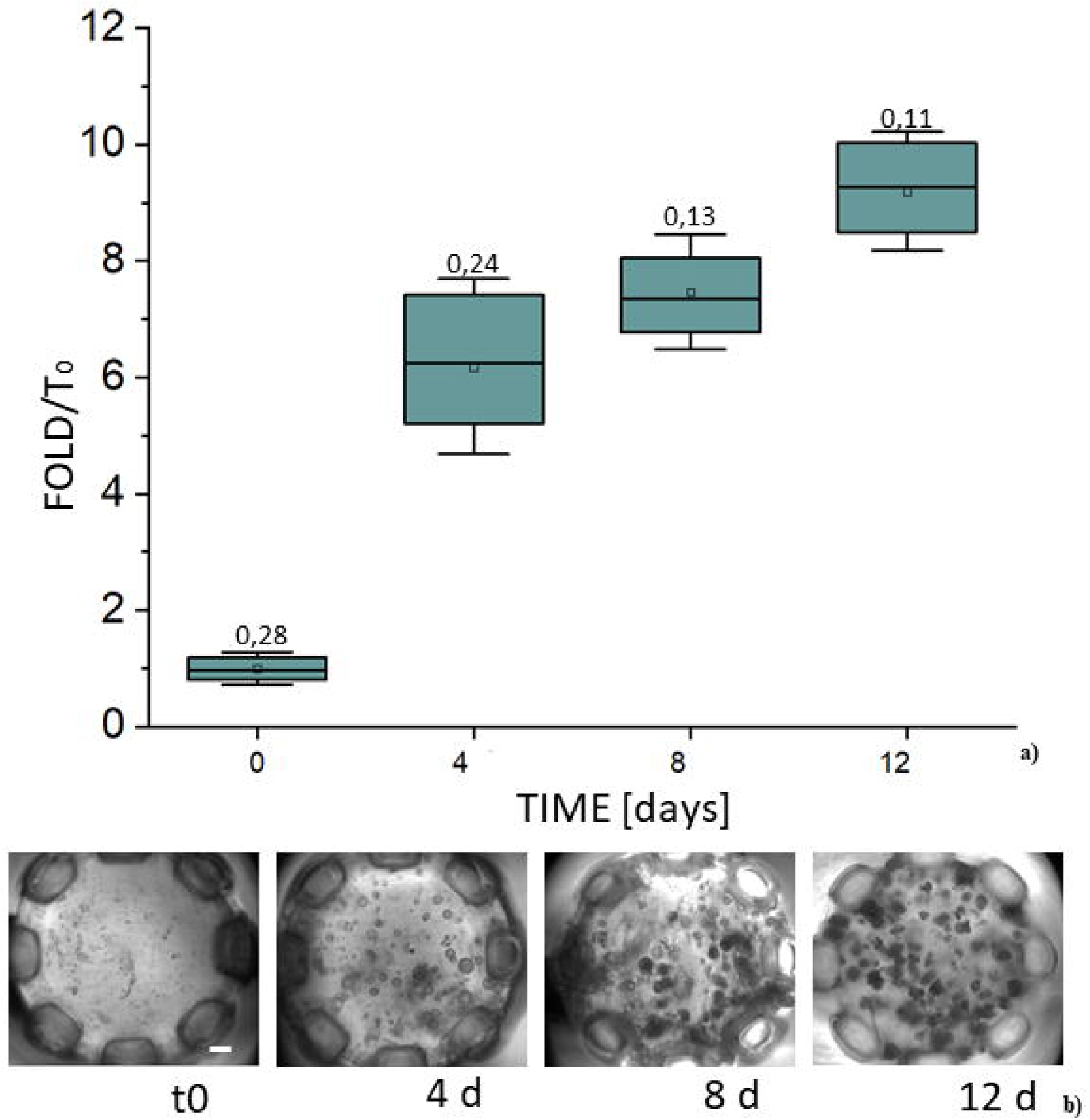
S**eeding and long-term culturing of Matrigel^TM^-embedded organoids.** PDO#17 suspended in 7mg/ml Matrigel was seeded and cultured in the NESTs system for 12 d. Every 96 hours, the viability of organoids was evaluated by CellTiter-Glo assay and brightfield imaging analysis. a) Growth rate (fold/t0) is reported as the ratio of the luminescence values at each time point relative to that at the time of seeding (t0). n= 36 replicates. Each box is defined by 25%-75% of the distribution ± SD (Standard Deviation). The square symbol represents the average, while the line represents the median value. The coefficient of variation CV is reported for each group. b) Representative brightfield images of the organoids at the reported time points. Scale bar: 200 µm.

### NESTs system allows homogenous growth of matrix-embedded PDOs

We subsequently evaluated the feasibility of culturing organoids in a 384-well plate format within the NESTs system. Three PDO lines, PDO#17, #61, and #63, were embedded in Matrigel^TM^ 5 mg/mL, seeded in the system, and cultured for 96 hours. The morphology and size of organoids grown in the NESTs system were compared with the same organoids grown in a conventional format (domes in a 24-well plate), using brightfield imaging analysis. Although PDOs showed the expected variability among lines (Fig. 3a and Fig. 2S), an ATP-dependent assay was performed to quantify the viability of organoids into the NESTs immediately after seeding (t0) and after 96 hours of culture. All the PDO lines exhibited an average growth of about 2 to 4-fold and data were consistent across independent samples at both time points (Fig. 3). Moreover, growth of the organoids in the NESTs system was comparable with that in conventional 96-well plates (Fig. 3c) and was unaffected by the concentration or the type of matrix used to embed PDOs (Fig. 4 and Fig. 1S). We then tested whether the NESTs system allows extended PDO cultures. PDO#17 was seeded in 7 mg/mL Matrigel^TM^. We chose this concentration of matrix because it is closer to that used for culturing organoids in conventional settings. Organoid viability was then assessed at different time points using an ATP-based assay (Fig. 4) and brightfield imaging analysis. The PDOs continued to grow over time (12 days), confirming that our system can support the homogeneous growth of matrix-embedded PDOs in a miniaturized format over extended periods of time. During this time, we did not observe any structural collapse of the droplets or matrix degradation.

### The NESTs system is suitable for drug screening on PDOs

In order to validate the NESTs system from a functional standpoint, we assessed its feasibility in registering responses to various drugs in a small-scale drug screening. Organoids derived from two different patients, PDO#63 and PDO#61, were seeded as described above and left untreated or treated for 96 hours with two drugs, bortezomib and olaparib. The luminescence values recorded at the end of the experiment using the ATP-based viability assay revealed that both PDO lines had been grown homogeneously (Fig. 5). As expected, proteasome inhibition by bortezomib resulted in sustained cell killing ^30^, whereas olaparib treatment elicited only modest effects. Altogether, these data support the potential to perform drug sensitivity tests in the NESTs system.

**Figure 5.**
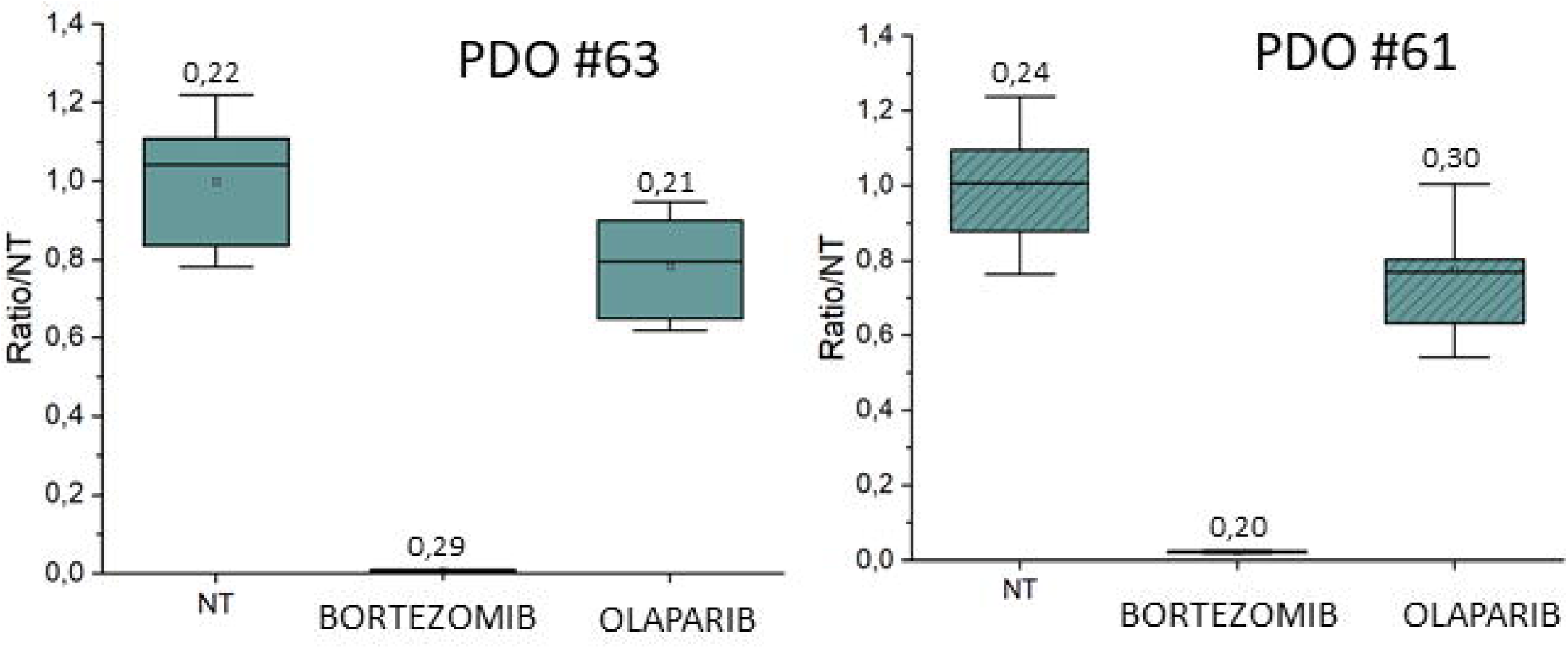
Drug sensitivity test on organoids seeded and cultured into the NESTs system. PDO#61 and PDO#63 were seeded and exposed to the indicated drugs. Viability of organoids was evaluated by CellTiter-Glo assay at 96 hours. n=12 replicates for each condition. Data are reported as the ratio of the luminescence values of the treated relative to the untreated samples (Ratio/NT). Each box is defined by 25%-75% of the distribution ± SD (Standard Deviation). The square symbol is the average, and the line is the median value. The coefficient of variation CV is reported for each group.

### The NESTs system supports imaging applications

Imaging-based screening, even in 3D cultures, is crucial for in-depth biological characterization. We explored the capability of using the NESTs system for imaging applications, in brightfield and confocal immunofluorescence (IF) microscopy for visualizing fluorescently labeled biological samples. Brightfield imaging analysis enabled the visualization of the organoid constructs throughout the entire depth of the matrix within the NESTs, which can be performed in the above-reported culture configuration, with the NESTs immersed in the wells of a clear-bottom 384-well plate (Figs. 4-6).

**Figure 6.**
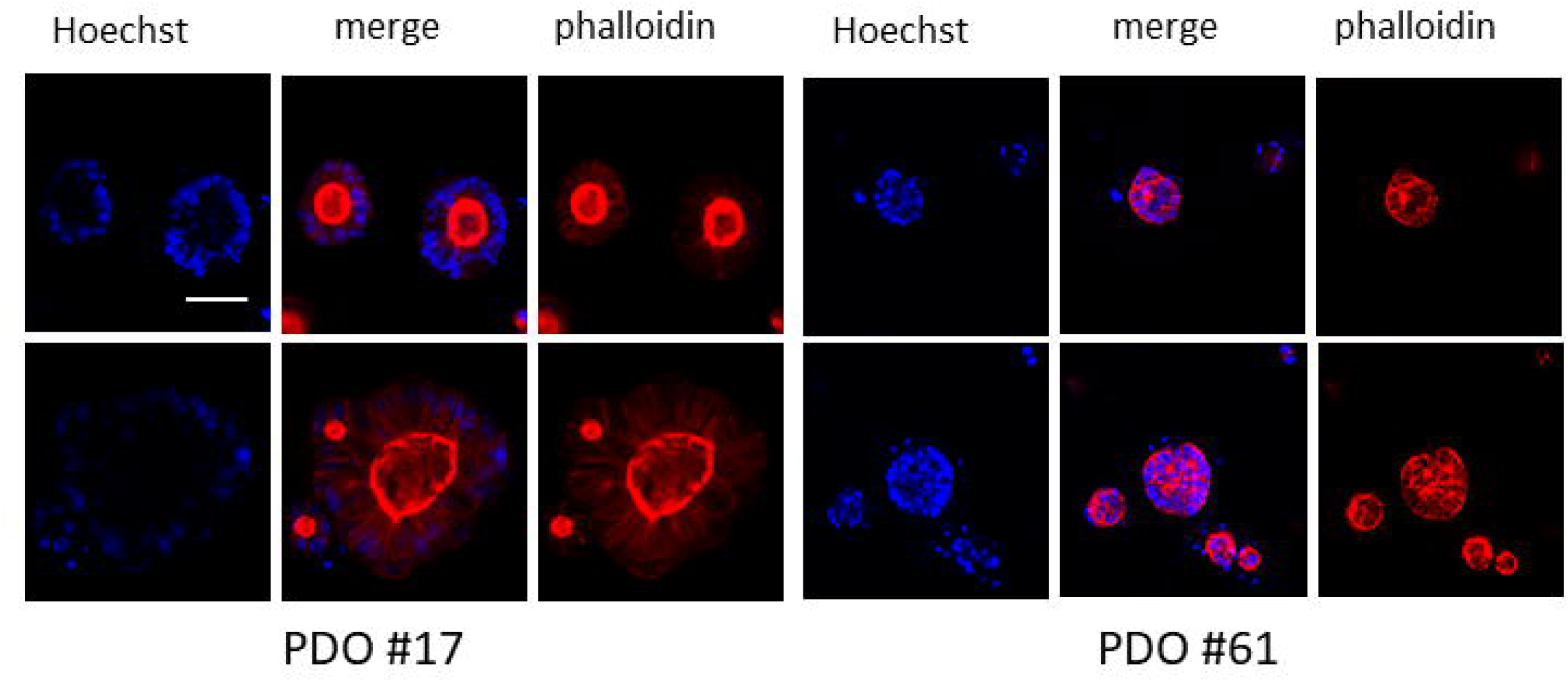
The NESTs system is suitable for imaging applications. PDO#17 and PDO#61 were seeded, cultured, fixed and stained in the NESTs system. Representative images reporting the staining of nuclei with Hoechst (blue) and the counterstaining of the cytosolic actin filaments with Rhodamine Phalloidin (red). Scale bar: 50 μm.

To allow the use of the system for advanced fluorescence-based confocal imaging, we employed a modified support to position the open side of the NESTs against the bottom of the well plate, minimizing the working distance. Using this dedicated culture setup, we validated the conditions for performing high-resolution confocal fluorescence imaging. Two PDO lines (PDO #17 and PDO #61) were seeded and grown within the system. Organoids were then fixed and stained with Hoechst, which binds specifically to the minor groove of DNA and stains the nucleus, and Phalloidin, which colocalizes with the F-actin component of the cytoskeleton (Fig. 6). Fluorescence analyses yielded consistent imaging of the structures distributed across a thickness of up to ∼300 µm (Fig. 6), consistent with the optical setup, and confirmed the possibility of performing imaging studies within the NESTs system.

## Discussions

In this study, we have described an innovative platform that combines a customized bioprinting system with modular 3D-printed components. This novel approach integrates basic microfluidic strategies and standard biological laboratory workflows, providing a scalable solution for the miniaturized culture of organoids. As a dispensing system we designed and validated a reusable and sterilizable cartridge by optimizing microfluidic strategies. When assembled with a syringe pump, the cartridge enables the precise deposition of homogeneous droplets of a biological matrix containing organoids at the microliter scale, ready for culture.

The configuration of the cartridge was designed to prevent the spontaneous sedimentation and a non-uniform distribution of the organoids across individual wells. The premature gelation of the biological matrix during the seeding is prevented by a refrigerated core integrated in the cartridge. The non-Newtonian pseudoplastic behavior of the sBME-based matrices, whose viscosity decreases as shear stress increases under flow conditions, is not compatible with pressure-driven extrusion systems. Thus, a syringe pump enabled accurate volumetric sample extrusion. To enable full sample loading into the cartridge and not in the syringe, where fluid velocity is minimal and sedimentation would occur, a buffer solution (PBS) was loaded in the syringe and separated from the sample by a small interposed air bubble whose volume is sufficient to maintain fluid separation without affecting the accuracy of the volumetric dispensing. Pump actuation resulted again in a volumetric extrusion of the sample. The configuration was compatible with the common needs of a sterile biological laboratory.

The standard methods used to describe the distribution and motion of the dispersed phase in multiphase flows (e.g., statistical description of multiphase dynamics) ^28^ were difficult to apply in this case, due to the size distribution of the organoids and the rheological properties of Matrigel^TM^. Therefore, the size and settings of the seeding components and protocols were defined by selecting a working point that compensated for particle migration, sedimentation, axial dislocation, and aggregate viability. This configuration was compatible with the common needs of a biological laboratory (sterility, a biosafety cabinet, and standard sizes of tools and instrumentation). The choice of a predominantly horizontal orientation of each spiral of the tubing, combined with the intermittent flow that enforces an almost horizontal flow direction, prevents sedimentation along the tubing length. A higher wall shear stress, compared to that in continuous flow at the same flow rate, was achieved thanks to the flatter velocity profile associated with the transient regime. Lower volume flow rates were ineffective in counteracting the sedimentation (Fig. 3S). The results obtained in this configuration were compared with preliminary experimental data (data not shown) obtained using a 90° tilted cartridge. In this configuration, non-uniform distribution and localized accumulation suggested sedimentation along the tube. Similar observations were made with a straight tube oriented vertically, where the organoids sedimented in the direction of flow. This resulted in their preferential accumulation in the initial droplets dispensed.

The cartridge consistently produced uniform droplets when bioprinting organoids, regardless of the matrix concentration in which they were embedded. In most of the experiments, organoid suspensions were prepared in a lightly diluted Matrigel^TM^ solution with a final protein concentration of 5 mg/mL (Figs. 3-7), chosen as a good compromise between the rheological properties of the matrix and the mechanical properties of the gelated drops^29^. However, we successfully seeded suspensions of organoids in higher Matrigel^TM^ concentrations (7 mg/mL), without protocol adjustments. To promote miniaturization and optimize reagent use in high-density 3D cultures, we successfully tested the seeding of 4, 6, and 8 μL drops, smaller than standard dome-shaped volumes in multiwell plates ^5,39^ (Fig. 2).

The NESTs system contains the 3D culture unit within the culture wells. We designed a structure mimicking microfluidic retention pins ^20^ to contain the matrix, while maintaining exposure to the culture medium and compartmentalization. Hanging the NESTs structures ^30^ upside down improves handling, preservation, and harvesting of the matrix sample, facilitating post-culture assays and allowing the units to be kept in culture for extended periods. The system features multiphase flow, with heterogeneous multicellular aggregates suspended in a non-Newtonian fluid. Pure Matrigel^TM^ undergoes a sol-gel transition at around 6 °C ^14^, which poses significant challenges for handling. To overcome this, it is typically mixed with other hydrogels or buffer solutions, such as PBS. This dilution may delay the attainment of gelation temperature, simplifying Matrigel^TM^ handling, and changes the final mechanical properties. In bioprinting applications, Matrigel^TM^ is often mixed with other polymers, such as alginate or gelatin ^13,31^, introducing additional biological variability ^32^. For drug screening applications, to simplify the seeding process and enable robotic handling ^33^, single cells/organoids are suspended in cell culture media containing low percentages (≤ 10%) of Matrigel^TM^ ^34,35^, a compromise that impacts organoid formation and branching due to the limited 3D scaffold support ^36,37^.

Our tool provides significant benefits in comparison to current microwell-based organoid culture systems. PDOs embedded in biological matrices can be seeded homogeneously and reproducibly into the final 3D culture units in a miniaturized and scalable system, eliminating the need for matrix removal or organoids growth on matrix-coated substrates ^33^. The miniaturization did not have a negative impact on the proliferation of organoids, whose growth rate in the NESTs system was comparable to that obtained in more conventional settings (Fig. 3), with an important reduction in the reagents usage (∼ 1/6). Moreover, our system supports short-term (96 hours) but also longer (12 days) culture of organoids (Figs. 4 and 6), with no significant matrix degradation during the experimental time frame. Remarkably, no selection of the initial population nor use of a specialized medium was performed to ensure the homogeneity and the reproducibility of organoids seeding and growth ^16^. The NESTs system brings benefits to the laboratory workflows: a high bioprinting rate of organoids droplets into the NESTs (36-unit to be prepared in approximately one minute) and a fast culture medium exchange by transferring the NESTs to a new plate. The NESTs configuration ensures the structural integrity of the culture units and enables matrix-embedded organoids to be extracted (centrifugation-based protocol) directly within the plate, making them available for post-culture characterization. The system provides benefits also compared to microfluidic approaches to organoids culture and screenings: microliter-scale droplets ^38,39^ suspended in sBME-based matrices within the culture medium, while useful for miniaturization, pose significant challenges in terms of individual manipulation and downstream processing. Similarly, microfluidic chips comprising interconnected compartments arranged side-by-side ^40^ do not allow for easy isolation or retrieval of the cultured units for further post-culture assays.

Finally, to feature the system for organoid-based HTS applications, we exposed organoids to bortezomib and olaparib, observing consistent effects across lines and replicates (Fig. 5), and validated the possibility of performing imaging studies (Fig. 6).

In summary, our system overcomes key limitations in using PDOs for HTS applications, advancing 3D culture models as standardized in vitro platforms. It combines volumetric bioprinting and 3D printing to support organoid growth in a miniaturized hanging-drop format, enabling precise seeding, consistent distribution, and streamlined workflows. Experimental validation confirmed its robustness and suitability for high-throughput drug screening and omics-based analyses of 3D cultures.

## Methods

### Dispensing system

The dispensing system includes a syringe pump that can dispense accurate and precise volumes of the sample (±1% accuracy), and a custom cartridge that is connected in series to the syringe and stores the entire sample to be dispensed for each assay. The dispensing system is designed to deposit individual microfluidic samples into dedicated 3D-printed structures, ensuring the homogeneity of organoids seeding, for culture and drug screening assays (Fig. 1). The cartridge consists of a PTFE tubing (BOLA^TM^, Bohlender GmbH, Grünsfeld, Germany), with Internal Diameter, ID, 1 mm and Outer Diameter, OD, 1,6 mm, coiled on a 3D printed support around an ice refillable compartment (a 15 mL centrifuge tube, Corning® Falcon®, Corning, NY, USA), to keep the bioink in a fluid phase, preventing premature gelation. The refrigerated coil is inserted into a plastic holder (a modified 50 ml centrifuge tube, Corning® Falcon®, Corning, NY, USA) that acts as a thermal insulator from the laminar air flow of the biosafety cabinet. The length of the coiled tube defines the maximum volume of the sample for each dispensing series. The cartridge used in these experiments can store up to 400 µL, and is designed for dispensing up to 300 µL. For most of its length, the tubing is positioned in a quasi-horizontal configuration (< 5°), and the flow inside the tubing (steps at 200 μL/min to dispense the drop, interspersed with 3 s of no flow) is designed to prevent sedimentation of organoid structures. The inlet of the cartridge is connected, using a custom 3D printed Luer-to-tube-fitting connector, to a glass syringe (5 mL Gastight TLL glass, Hamilton, Reno, NV, USA) actuated by a syringe pump (PhD2000, Harvard Apparatus, Holliston, MA, USA). The syringe is filled with a buffer solution such as PBS (Phosphate Buffered Saline, Sigma-Aldrich, St. Louis, MI, USA). A small air bubble (∼ 10 – 50 μL) created on purpose during the cartridge filling is left between the buffer fluid and the sample to avoid mixing and the uncontrolled dilution of the sample by PBS. The outlet of the cartridge is connected to another short tubing segment (1 cm length, ID 0,5 mm, OD 1,6 mm), allowing the proper formation of drops and their release to the target 3D printed structures, referred to as NESTs in the following text. The dispensed drops are deposited in a series of 4, 6, and 8 μL to fill the respective NESTs (Fig.3).

### 3D printing of system components

The fabrication of structures and tools was performed by stereolithography (SLA) 3D printing (Formlabs® Form 3B, Formlabs, Somerville, MA, USA). Structures are printed using a biocompatible resin, Biomed Clear V1 (Formlabs®, Somerville, MA, USA), certified for long-term skin/mucosal membrane contact ^41^. A standard post-processing protocol is applied, consisting of a washing step in isopropyl alcohol of 20 min (IPA, 415158 Carlo Erba, Dasit Group, Milano, IT) and an UV curing step (Form Cure, Formlabs®, Somerville, MA, USA) at 60 °C for 60 min, followed by immersion in deionized water for additional rinsing steps, drying under a laminar air-flow hood, and final sterilization by UV exposure in sealed Petri dishes for 20 min. Additional components, including tools for handling and transferring organoid supports during gelation, a multiwell plate holding frame, fluidic connectors, and mechanical components of the cartridge, were fabricated using the same 3D printing technology and material.

### 3D printed culture unit supports

Each NEST is a 3D-printed structure designed to accommodate a microliter drop of a biological matrix, creating the 3D-culture unit. Structures are characterized by hydrophilic surface properties (contact angle ∼50°), enabling the sample to fill the cavities. We prototyped different structures able to host a maximum volume of 4, 6, and 8 µL (Fig. 1). The NEST is a pseudo-cylindrical structure defined by a circular arrangement of pins that enclose an internal cavity open at the top. The pins alternate with open fenestrations (OD ∼2,4 mm and ID ∼1,9 mm), which allow lateral contact between the matrix and the cell culture medium. The different heights of the pins correspond to the maximum sample volumes (1,25, 1,9 or 2,5 mm, respectively) that can be contained inside the cavity. When the NEST is immersed in the culture medium, around 40% of the lateral surface of the matrix comes into contact with the cell culture medium and is available for diffusional exchange. The NESTs are positioned at the top of pillars arranged in 36-unit arrays (6 × 6) on custom-designed supports. The pillars are dimensioned to fit within the layout of a standard 384-well plate, allowing each NEST to be individually immersed in a single well while maintaining the culture units suspended within the culture medium. In the culture configuration, the open surface of each NEST is located 3 mm above the bottom of the well.

### Organoids samples

Sub-confluent PDOs were harvested, mechanically and enzymatically dissociated into single cells by incubating with TrypLE Select enzyme (Gibco, 10272625) ^42^. Cells were embedded in the matrix and seeded in 100 µL-domes in a 24-well flat-bottom cell culture plate (Corning). Domes were overlaid with complete medium ^42^, allowing the generation of small PDOs for 2-3 days (characteristic size ≤ 50 µm), and then recovered from the matrix with the Cell Recovery solution (Corning, 354253). Recovered PDOs were diluted in solutions of growth factor-reduced.

Matrigel™ (Corning, 356231) or Cultrex™ RGF Basement Membrane Extract type 2 (R&D Systems, 3533-010-02) in PBS. Depending on the batch of the matrices, the solutions were diluted to fit a concentration of proteins of ∼5 mg/mL (generally obtained in solution with 50% of Matrigel™). In culture settings employing Cultrex™, the matrix was prepared to achieve a protein concentration of 7 mg/mL (generally obtained in solution with 70% solution of Cultrex™). The suspensions were set to obtain a cell density of 375.000/ml, corresponding to approximately 3.000 cells aggregated in organoids within each 8 µL NEST unit. For long-term cultures, organoids were suspended in a 7 mg/mL Matrigel^TM^ solution, and 1.500 cells aggregated in organoids were seeded in each NEST unit. For culturing in 96-well plates (Corning #3917), 50 µL of the organoid suspensions were seeded in each well and overlaid with 100 µL of medium. Organoids used in this study were derived according to established protocols ^9^ from patients undergoing surgical resections of colorectal cancer metastases to the liver, and belong to a biobank of genomically and clinically annotated organoids (manuscript in preparation). The protocol for the use of human samples was approved by the Institutional Review Board (Ethic Committee IRCCS Ospedale San Raffaele, Milan, Italy; protocol ACC_ORG) in compliance with the principles of the Declaration of Helsinki, and all participants in the study signed informed consent.

### Seeding protocol in the NEST units

The seeding protocol (Fig. 1) for generating the organoid culture units comprises three steps: (i) bioprinting of the matrix drop into the NESTs structures, (ii) matrix gelation, and (iii) positioning of the filled NEST into the culture wells containing medium. The dispensing system is placed under the biosafety cabinet. To prepare the system for the seeding, the sample (organoids suspended in biological matrix) must be preloaded into the cartridge. The sample is manually transferred by a 1 mL syringe (Terumo Corporation, Japan) through the inlet connector (∼ 200 µL/s). Once the sample is preloaded, a microliter air bubble is released from the same syringe and left in the cartridge tube to separate PBS from the sample. The connector is then locked to a glass syringe, which is activated by a syringe pump. The pump is actuated at 200 µL/min, bringing the sample flowing through the cartridge to the outlet. Once the cartridge has been filled, 60 µL of the sample is extruded, but not seeded in order to avoid any inhomogeneity related to the filling step (data not shown). Finally, the pump is actuated to extrude 4, 6, or 8 µL. During the seeding, each of the 36-unit NESTs, anchored in a 50 mm Petri dish inside a 3D-printed holder, each NEST is manually aligned beneath the cartridge outlet tube. The drops are then collected one by one in the NESTs. The drop fills the internal volume of the NEST, led by gravity, capillarity, and surface tension, and settles in contact with the air in the open spaces between the pins. The self-filling property of the NEST enables compensation for minor misalignments during droplet deposition, ensuring consistent cavity filling and enhancing reproducibility. Once the 36-unit NESTs are filled, the matrix gelation process is carried out by flipping the set upside down and suspending it on the longer integrated alignment pillars. The set is then stored inside a closed polystyrene Petri dish in an incubator at 37 °C with 5% CO_2_ for 10 min. In the third and last step, the 36-unit is placed in culture in a 384-well plate (flat clear bottom – squared – 112 μL total volume, Corning, #3765). The layout of the 36-unit ensures that each NEST is aligned with a single well (Fig. 1n), side-by-side, dipped in 70 µL of complete organoid medium. The multiwell plate is then covered with a lid, secured in place by a 3D-printed adapter, before being placed in culture for the desired time.

### Viability test

An ATP-based assay was used to measure cell viability, which then served as an indicator of the homogeneity of organoids seeding and growth in different NESTs and to quantify the response to drugs in drug screening experiments. The units of matrix containing the organoids were dislodged from the chip and directly transferred into a new empty 384-well white plate (Corning, #3765) by centrifugation (3 min at 800 g). Then, 70 µL of a 1:1 mix of CellTiter-Glo 3D (Promega, G9681) and Advanced DMEM/F12 (Gibco, 12634028) was added to each well. After 45 min of incubation at room temperature, according to the manufacturer’s instructions, the luminescence signal was measured on a Mithras LB 940 microplate reader (Berthold Technologies) and reported as Relative Luminescence Units (R.L.U.). The mean of the luminescence values of independent samples, calculated for each condition and time point, was reported as mean ± standard deviation (SD). The number of replicates for each experiment is indicated in the corresponding figure legend. Luminescence values were normalized with respect to those at seeding time and non-treated (NT) for growth rate evaluation and drug screening experiments, respectively. Viability assays in 96-well plate were performed by adding in each well 50 µL of CellTiter-Glo 3D.

### Drug screening on PDOs grown and treated in NEST units

Bortezomib at 1 µM and olaparib at 10 µM (both from MedChem Express, Monmouth Junction, NJ, USA) were used for drug screening experiments. Organoids were seeded as described above and exposed to drugs. From a stock concentration of 1000X in the vehicle, drugs were further diluted to 1X final concentration in complete organoids medium, and 70 μL of this solution was added to each well. DMSO-treated organoids served as control. DMSO concentration in the medium was kept under 0.1%. Organoids were incubated with the compounds at 37 °C for 96 h with no further changes of media or compound refresh. Then the cultures were subjected to viability assays as described above. The mean of the luminescence values of technical replicates was calculated for each condition. Results are reported as the mean percentage of viable cells in drug-treated relative to DMSO-treated control samples.

### Long culture of organoids in the NEST units

Organoids were plated as described above and cultured for up to 288 h (12 days). The medium was renewed every 96 h (4 days) by removing the support from the used multiwell plate and placing it into a new plate containing fresh medium. The growth of organoids was evaluated at intervals of 96 h by CellTiter-Glo assay and bright-field imaging (Axio Observer inverted microscope, ZEISS).

### Imaging

Imaging of the organoids cultured either in the NESTs supports or in the regular multiwell plates was performed using a brightfield microscope (Axio Observer inverted microscope, ZEISS). For confocal imaging, organoids were plated as described above on NESTs grid supports specifically designed for imaging applications. This modified NEST holder allows the open surface of the NESTs to be positioned closer to the bottom of the plate. This reduces the working distance with the microscope objective. Organoids were stained directly into the NESTs, as already described ^29^. Briefly, PDOs were fixed in a solution of 4% paraformaldehyde (Electron Microscopy Sciences, 157-8) and 2% sucrose (Sigma-Aldrich, S0389) for 30 minutes and then permeabilized with 0.1% Triton X-100 (Sigma-Aldrich, T8787) for 8 minutes. Nonspecific antigens were blocked in a solution of 5% BSA (Sigma-Aldrich, A8022) and 10% goat serum (Sigma-Aldrich, G9023) for 30 min, and then organoids were incubated for 2 h with a solution of 10 μg/mL Hoechst 33342 (Invitrogen, H3570) and 4 μg/mL rhodamine phalloidin (Invitrogen, R415) in 2.5% BSA. At the end of the staining, the NESTs 36-unit was placed on a new 384-well black glass-bottom microplate (Ibidi, #88407) filled with PBS and stored at 4 °C before imaging. 16 bit -1024x1024 pixels images were acquired using an Olympus Fluoview 3000RS confocal laser scanning microscope with an oil 30X/1.05NA objective.

## Funding

This work was supported by a number of Funding Agencies:

The Accelerator Award A26815 ‘Single-cell cancer evolution in the clinic’, funded through a partnership between Cancer Research UK (London) and Fondazione Italiana per la Ricerca sul Cancro (AIRC, Milan) (to G.T. and G.D.)

The Italian Ministry of University and Research, PRIN Project ’Microfluidic Platform for Cancer Drug Screening – MicroCaDS’, The European Union - Next Generation EU, NRRP (National Recovery and Resilience Plan) Mission 4, Component 1, CUP D53D23003310006 (to G.D. and G.T.)

‘Multilayered Urban Sustainability Action - MUSA’, The European Union – Next Generation EU, NRRP, Mission 4, Component 2 (to G.D.)

The Italian Ministry of Health, grant RF-2021–12374586 (to G.T.)

The Italian Ministry of University and Research, Project PNC0000001 ‘D34 Health—Digital Driven Diagnostics, prognostics and therapeutics for sustainable Health care’, CUP B53C22006090001 (to G.T.)

The Italian Ministry of Health, Project PNRR-MCNT1-2023-12378347 ‘Patient-derived organoids microfludic screenings to define treatments for metastatic colon cancer’, The European Union - Next Generation EU, NRRP Mission 6, Component 2, CUP C43C24000220007 (to L.A. and G.T.).

## Declarations

### Human tissue samples

All procedures were approved by the Institutional Review Board of the IRCCS San Raffaele Scientific Institute and were conducted in compliance with all relevant ethical regulations. Patient samples were collected at the Clinical Department at San Raffaele Hospital (Milan, Italy) under written informed consent in agreement with the Declaration of Helsinki (Protocol “ACC_ORG”, Milan, Italy). All data were stored according to best practices, protecting patient confidentiality and data integrity

## Acknowledgements

The authors wish to thank all the members of the Dubini and Tonon labs for helpful discussions and suggestions, alongside the clinical colleagues of the Aldrighetti team; Francesca Letizia Capelli for introducing and supporting us in the SLA 3D printing; Monica Piergiovanni for brainstorming on the design of the system; Cesare Covino and Valeria Berno at the Advanced Light and Electron Microscopy BioImaging Center (ALEMBIC) core facility of IRCCS San Raffaele Scientific Institute for expertise and assisting with instrumentation; Cinzia Felicita Sala for the generation of the Core Genome Platform (C.G.P.) framework.

## Author contributions

E.B. Conceptualization, Supervision, Methodology, Investigation, Data curation, Writing_original draft. O.A.B. Conceptualization, Supervision, Methodology, Investigation, Data curation, Writing_review and editing. P.D.S. Methodology, Writing_review and editing. G.F.G, C.F. and J.M.B Investigation, Data curation. Writing_review and editing G.G. Investigation. F.R. and L.A Conceptualization, Resources. R.K Conceptualization, G.T. Conceptualization, Funding acquisition, Supervision Project administration, Resources, Writing_review and editing. G.D. Conceptualization, Funding acquisition, Supervision, Project administration, Resources, Writing_review and editing.

## Supplementary Figures

**Figure 1S:** Seeding and culturing of matrix-embedded organoids in the NEST system are unaffected by the concentration or the type of matrix used. PDO#63 suspended in 7mg/mL Cultrex^TM^ was seeded and cultured for 96 hours in the NESTs system ATP content was measured by CellTiter-Glo assay at the time of seeding (t0) and at the end of culture. Growth rate (fold/t0) is reported as the ratio of the luminescence values at each time point relative to that at the time of seeding (t0). n= 18 at t0, n= 36 at 96h. Each box is defined by 25%-75% of the distribution ± SD (Standard Deviation). The square symbol represents the average, and the line represents the median value. The coefficient of variation CV is reported for each group

**Figure 2S:** PDOs cultured in the NESTs system grow as in conventional conditions. Representative brightfield images (5X objective lens) of three PDO lines seeded and cultured in the NESTs system (a and b), or in conventional conditions (c and d). a) and d), scale bar: 100 μm) , scale bar: 200 μm,. a) and d) are zoomed frames of b) and c) respectively.

**Figure 3S:** Homogeneity of seeding procedure into the NESTs system depends on the flow rate. Drops of matrix embedded organoids were seeded at two different flow rates and ATP content was evaluated by CellTiter-Glo assay. Data are reported as luminescence values (R.L.U.) The drops seeded at 200 µL/min showed values. within the expected range (red dotted lines), while the drops seeded at 25 µL/min showed lower, more dispersed values. Each box is defined by 25%-75% of the distribution ± SD (Standard Deviation). The coefficient of variation CV is reported for each group. Scale bar 200 µm.

