## Supplementary figures and images for "3D printing and bioprinting for miniaturized and scalable hanging-drop organoids culture"

### supplementary Fig. 1

FIG_1_SUPP

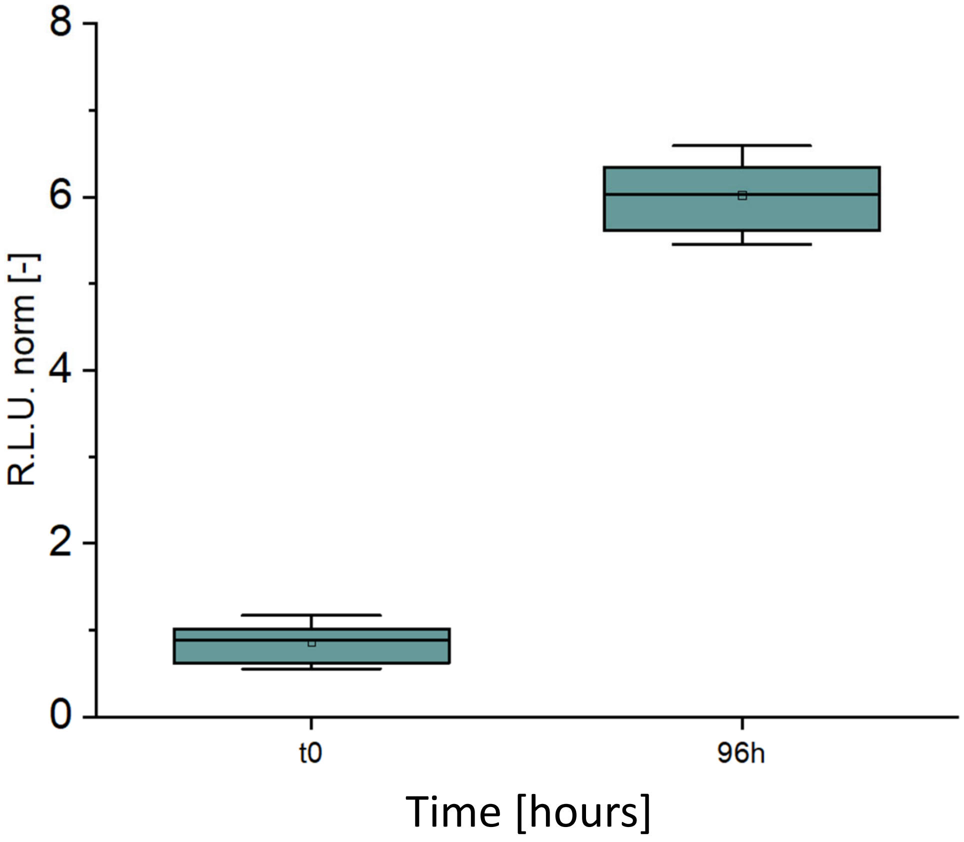

### supplementary Fig. 2

FIG_2_SUPP

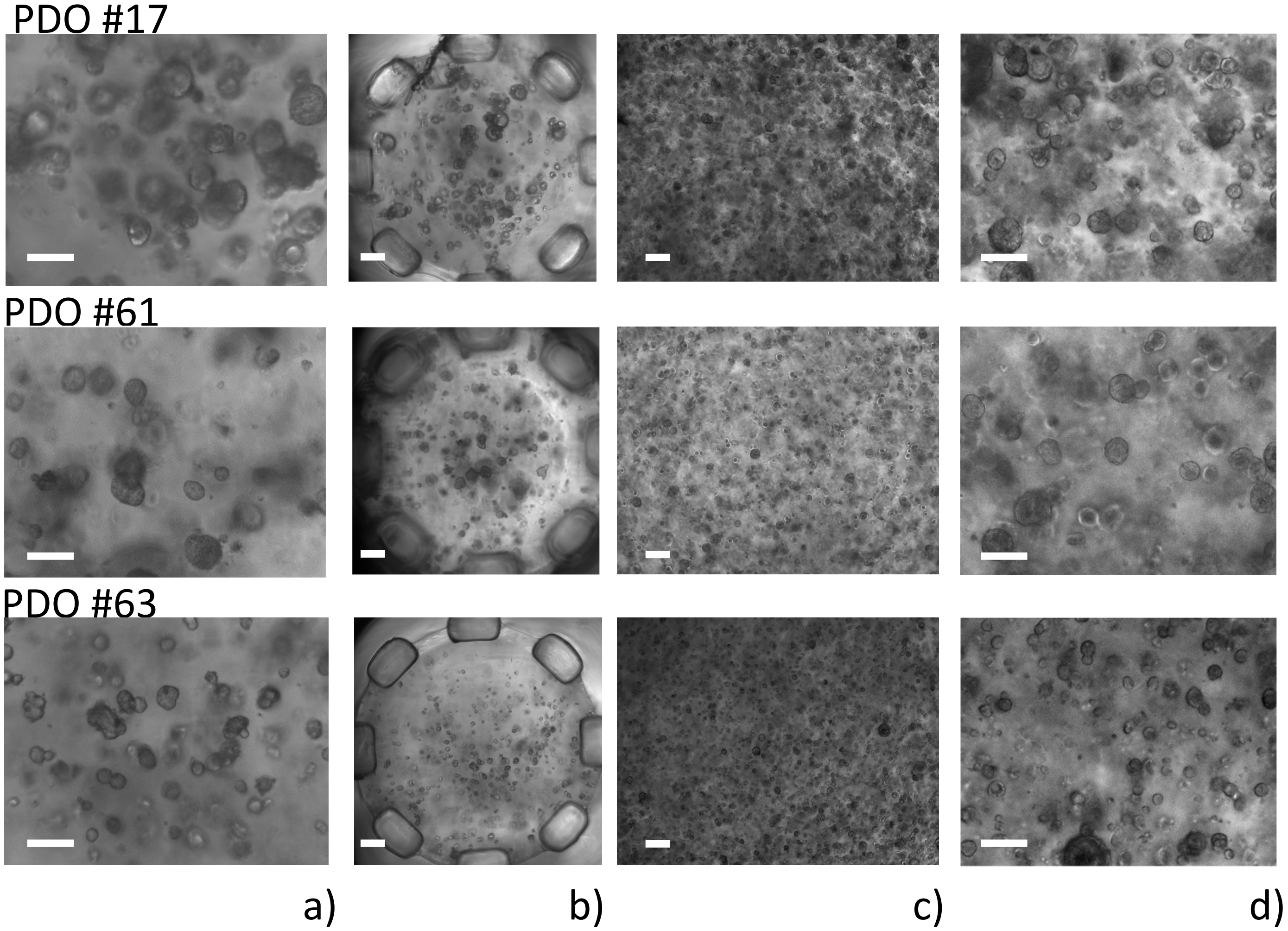

### supplementary Fig. 3

FIG_3_SUPP

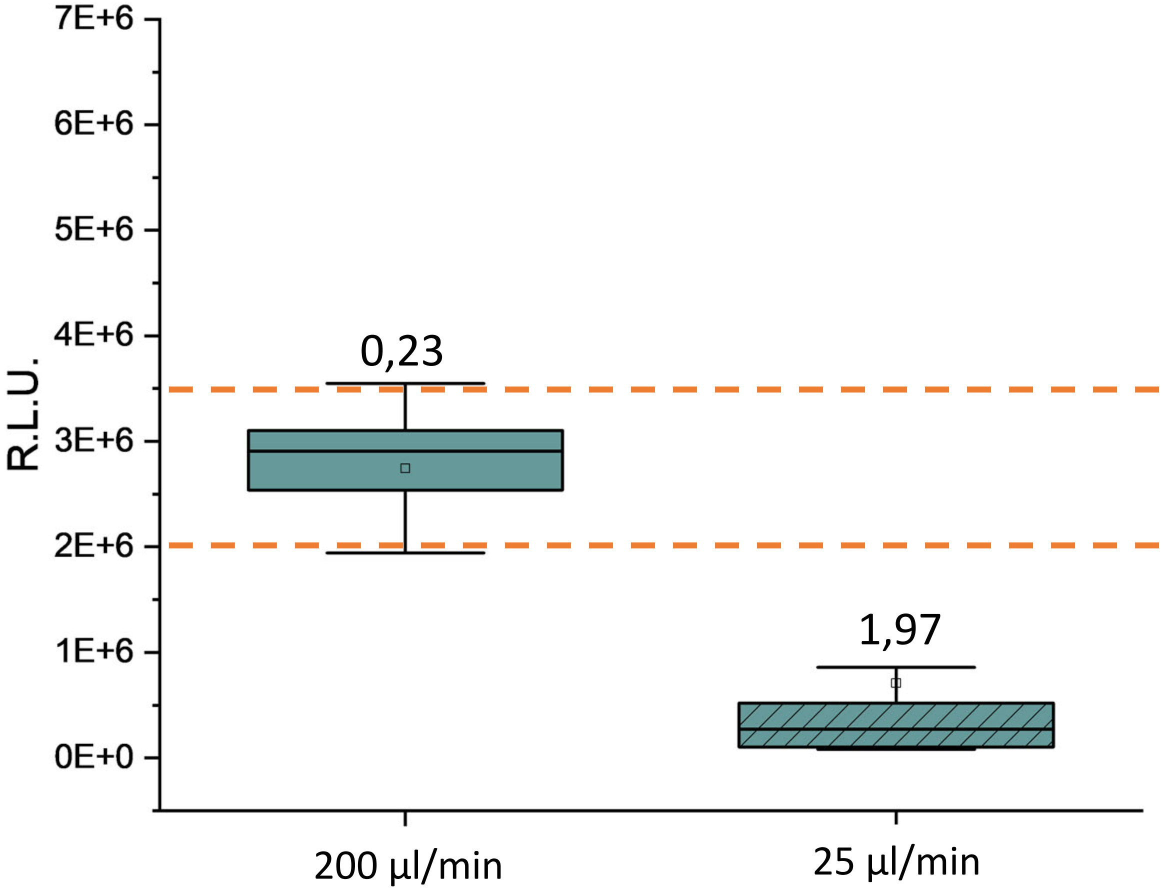
